# Proliferation of HIV-infected renal epithelial cells following virus acquisition from infected macrophages

**DOI:** 10.1101/2020.03.27.011916

**Authors:** Kelly Hughes, Guray Akturk, Sacha Gnjatic, Benjamin Chen, Mary Klotman, Maria Blasi

## Abstract

**Objectives:** HIV-1 can infect and persist in different organs and tissues, resulting in the generation of multiple viral compartments and reservoirs. Increasing evidence supports the kidney as such a reservoir. Previous work demonstrated that HIV-1 infected CD4+ T-cells transfer virus to renal tubule epithelial (RTE) cells through cell-to-cell contact. In addition to CD4+ T-cells, macrophages represent the other major target of HIV-1. Renal macrophages induce and regulate inflammatory responses and are critical to homeostatic regulation of the kidney environment. Combined with their ability to harbor virus, macrophages may also play an important role in the spread of HIV-1 infection in the kidney.

**Design and Methods:** Multiparametric histochemistry analysis was performed on kidney biopsies from individuals with HIV-1 associated nephropathy (HIVAN). Primary monocyte-derived macrophages were infected with a (GFP)-expressing replication competent HIV-1. HIV-1 transfer from macrophages to RTE cells was carried out in a co-culture system and evaluated by fluorescence-microscopy and flow-cytometry. Live imaging was performed to assess the fate of HIV-1 infected RTE cells over time.

**Results:** We show that macrophages are abundantly present in the renal inflammatory infiltrate of individuals with HIVAN. We observed contact-dependent HIV-1 transfer from infected macrophages to both primary and immortalized renal cells. Live imaging of HIV-1 infected RTE cells revealed four different fates: proliferation, hypertrophy, latency and cell death.

**Conclusions:** Our study suggests that macrophages may play a role in the dissemination of HIV-1 in the kidney and that proliferation of infected renal cells may contribute to HIV-1 persistence in this compartment.

## INTRODUCTION

HIV-1 persists indefinitely in infected individuals despite suppression of HIV-1 replication with antiretroviral therapy (ART)^[1]^. The persistence of HIV-1 in different sites throughout the body of infected individuals, including lymph-nodes, gut, liver, central nervous system and kidneys^[2-4]^, has important implications for viral pathogenesis and cure strategies. HIV-1 infection is associated with end-stage renal disease (ESRD), especially in African American individuals^[5]^. The association between HIV-1 infection of kidney epithelial cells and renal pathology was first demonstrated in 1992 using a transgenic mouse model^[6]^ that recapitulated HIV-associated nephropathy (HIVAN)^[7]^. Furthermore, HIV-1 nucleic acids have been detected in human renal biopsies^[8-10]^ and sequence analysis provided strong evidence that the kidney is a separate compartment from the blood for HIV-1 replication^[11]^. While widespread use of ART has significantly decreased the incidence of HIVAN, ESRD remains common among HIV-1 positive individuals^[12]^.

We have previously shown that HIV-1 DNA and RNA persist in renal epithelial cells despite treatment with ART^[9]^. It is unclear whether persistent virus in the kidneys contributes to chronic inflammation and renal pathology. Additionally, a study conducted on HIV-1 positive individuals that received kidney transplants from HIV-1 negative donors, demonstrated the presence of viral nucleic acid in the allografted kidney epithelial cells, despite undetectable plasma viremia^[13]^. Because renal epithelial cells lack both the CD4 receptor and the CCR5, CXCR4 coreceptors required for cell free virus infection^[14, 15]^, a potential route of infection of the allografted kidney is the formation of a virological synapse between recipient infected T-cells and donor kidney epithelial cells, as we have previously shown *in vitro*^[16, 17]^.

Macrophages, also a major target for HIV-1, are abundant in inflamed tissues. Compared to T-cells, virus replication is slower in macrophages and macrophages are more resistant to the cytopathic effects of HIV-1 infection^[18]^. It has been shown that infected macrophages can avoid the cytopathic effects of viral budding by storing newly produced viral particles in membrane pockets^[19, 20]^, which allows tissue-resident macrophages to survive for prolonged periods. Given the important roles of macrophages in kidney homeostasis and in the response to acute and chronic kidney injury^[21, 22]^, we hypothesized that HIV-1 infected macrophages could contribute to initiating and maintaining infection of renal epithelial cells.

Here, we demonstrated that macrophages transfer virus to renal tubule epithelial (RTE) cells through direct contact. Once infected RTE cells can in turn mediate infection of monocytes and T-cells, supporting a “ping-pong” infection model between immune cells and epithelial cells that sustains HIV-1 infection within the kidney. Live imaging of flow-sorted HIV-1 infected renal cells revealed the downstream consequences of RTE cells infection. Some cells undergo cell-death or hypertrophy that could account for the renal injury associated with HIV-1 infection. Other infected cells undergo multiple rounds of cell division, with or without transcriptional silencing. Our study highlights the mechanisms of HIV-1 spread and persistence in the kidney.

## METHODS

### Multiplexed Immunohistochemistry on Renal Biopsies

We conducted a retrospective, immunohistochemical analysis to characterize the inflammatory infiltrate in confirmed HIVAN cases on archived formalin fixed paraffin embedded (FFPE) renal biopsies. The multiplexed immunohistochemical consecutive staining on single slide (MICSSS) approach was employed as previously described^[23, 24]^. We examined 5 HIVAN renal biopsies by MICSSS to quantify T-cells (CD3), CD8+ T-cells (CD8), neutrophils (CD66b), and monocyte/macrophages (CD68). Whole slide images (WSI) of the mentioned markers were analyzed by using QuPath, an open source image analysis platform^[25]^. Biopsy sections on the WSIs were fully annotated and quantification of positive cells on these biopsies was performed by setting the color vectors for hematoxylin and 3-Amino-9-ethylcarbazole (AEC) substrate, nuclear segmentation of cells in the annotation area and random forest-based classification of positive cells, respectively^[24]^ Staining for one out of five HIVAN diagnosed biopsies is shown in **figure 1**.

**Figure 1.**
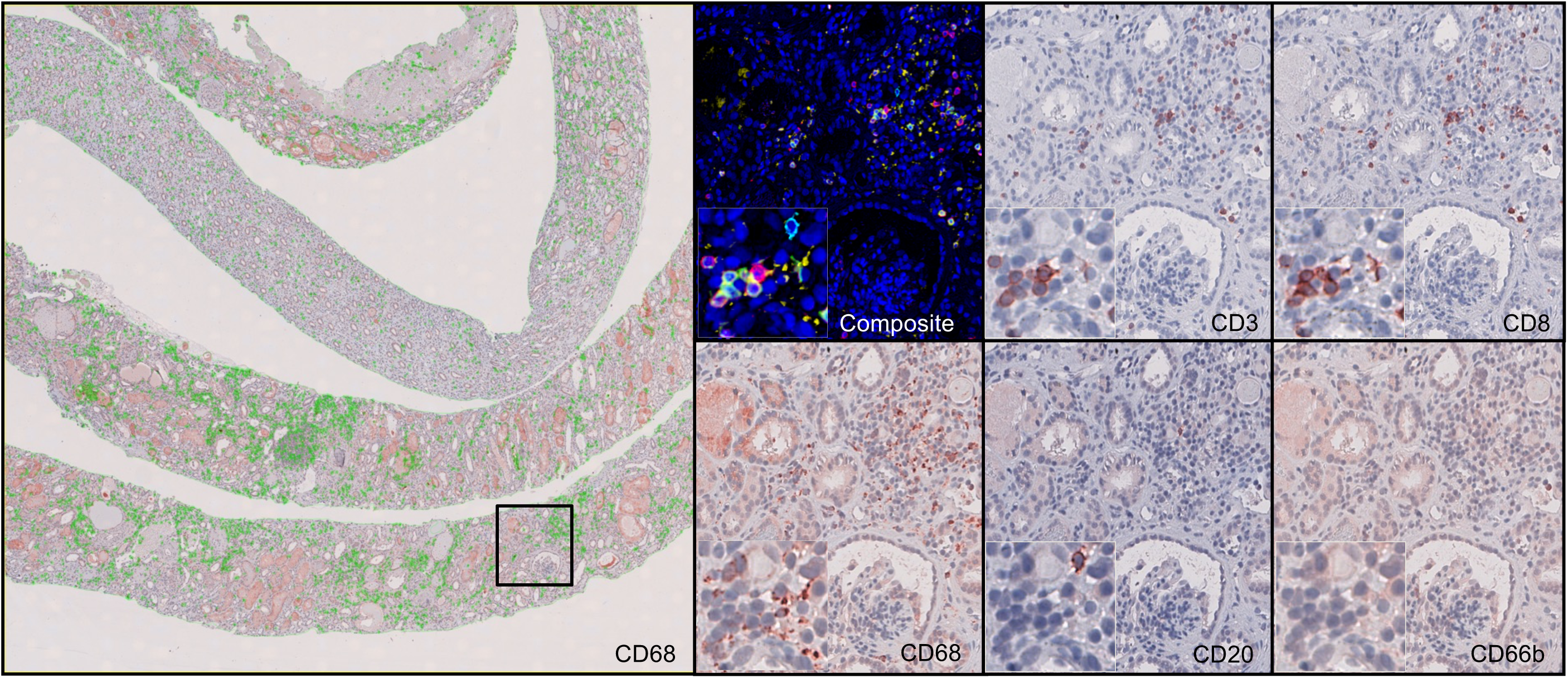
Presence of macrophages in interstitial infiltrates observed in HIVAN. Multiplexed immunohistochemical consecutive staining on single slide (MICSSS) analysis on a kidney biopsy from a HIV-1 positive subject with HIVAN. The mononuclear infiltrate was characterized by serially staining the tissue with markers for macrophages (CD68), T-cells (CD3), cytotoxic T-cells (CD8), B cells (CD20) and granulocytes (CD66b). Composite figure is produced by using each marker image with image registration, color inversion and image overlay method. Composite figure shows CD3+ cells in red, CD8 positive cells in green, CD20 positive cells in cyan, CD66b+ cells in magenta, and CD68 positive cells in yellow color. The shown staining of interstitial inflammatory infiltrates is from one representative HIVAN renal biopsy from a formalin fixed paraffin embedded tissue sample. The number of positive cells/mm2 for each of the analyzed marker is as follow: 499 for CD20, 1102 for CD3, 998 for CD66b, 208 for CD68 and 544 for CD8.

### Primary Cells and Cell Lines

HEK 293T Lenti-X cells (Clontech Laboratories, Mountain View, CA) were maintained in Dulbecco’s Modified Eagles medium (Thermo Fisher Scientific, Waltham, MA) supplemented with 10% fetal bovine serum (GE Healthcare Life Sciences, HyClone Laboratories, South Logan, UT) and 100 units per ml of penicillin– streptomycin–glutamine (PSG) (Thermo Fisher Scientific). The previously described human proximal tubular epithelial cell line HPT-1b^[26]^ was maintained in renal epithelial cell growth media (Lonza, cat. no. CC-4127). To generate the HPT-1b-mCherry cell line, constitutively expressing the mCherry gene, HPT-1b cells were transduced with 1 multiplicity of infection (MOI) of the SIV-based lentiviral vector^[27]^ expressing both the mCherry and neomycin resistance gene under the CMV promoter (GAE-CMV-mCh-IRES-Neo). Stably transduced cells were selected by treatment with 800 µg/mL of G418 sulfate (Corning) for 2 weeks.

CD14+ monocytes were isolated from healthy donor (age range 19-30 years old, 2 Caucasian males and 1 Black African American female) peripheral blood mononuclear cells (PBMC) by magnetic bead separation (MACS cell separation; Miltenyi Biotech, Cologne, Germany) and differentiated into macrophages by culture for 7 days in RPMI1640 medium supplemented with 10% fetal bovine serum (GE Healthcare Life Sciences, HyClone Laboratories, South Logan, UT) and 100 units per ml of penicillin– streptomycin (Thermo Fisher Scientific) and 20 ng/mL of MCSF (Peprotech, #AF-300-25, Rocky Hill, NJ, USA). Fresh medium containing MCSF was added every 2-3 days. Allogeneic primary renal tubule epithelial cells were obtained by culturing urine derived cells as previously described^[28, 29]^. The research protocol was approved by the Duke University institutional review boards (Pro0040696) and informed consent was obtained. The monocytic cell line THP-1 (ATCC, Cat.# TIB-202) and the CEM-SS T-cell line^[16]^ were maintained in RPMI1640 medium supplemented with 10% fetal bovine serum and 100 units per ml of penicillin–streptomycin.

### HIV-1 production and titration

The proviral plasmids pNL4.3-ADA-GFP^[30]^ and pSF162-R3Nef+^[31]^ were generously provided by Dr. Eric Cohen (Montreal Clinical Research Institute) and Dr. Amanda Brown (Johns Hopkins University), respectively. The pNLGI-JRFL and pNLGI-Δenv proviral plasmids have been previously described^[32, 33]^. For production of HIV-1 infectious molecular clones, 3.5×10^6^ Lenti-X cells were transfected with 10 µg of the HIV-1 plasmid using the JetPrime transfection kit (Polyplus Transfection Illkirch, France) following the manufacturer’s recommendations. To allow entry into macrophages the envelope-mutant virus (NLGI-Δenv) was pseudotyped with the JRFL env glycoprotein by co-transfection of pNLGI-Δenv with pJRFL^[34]^ at a ratio of 4:1 (10 ug total). At 48- and 72-hours post-transfection, culture supernatants were concentrated by ultracentrifugation for 2 hours at 23.000 RPM on a 20% sucrose cushion. Pelleted viral particles were resuspended in 1× phosphate-buffered saline (PBS) and stored at −80 °C until further use. Each viral stock was titered using the GHOST(3)CXCR4+CCR5+ reporter cell line (NIH AIDS Reagent Program (cat# 3942))^[35]^.

### Macrophage Infection and RTE Co-culture

Macrophages were infected at a MOI of 2.5, or 6 of HIV-ADA, HIV-SF162 or HIV-JRFL for 8 hours in serum-free RPMI medium, followed by a return to culture in complete RMPI media for 7 days. The percentage of infected cells was quantified by assessing GFP expression by flow cytometry. HPT-1b-mCherry cells or primary urine-derived RTE cells, stained with 30uM of the CellTrackerDeepRed (DR) fluorescent dye (Molecular Probes, Eugene, OR, USA; cat# C34565), were co-cultured with infected macrophages at a 1:4 ratio for 6 days in complete RPMI medium in presence or absence of a transwell membrane to block cell-cell contact. Co-cultures wells were then analyzed for the presence of mCherry and GFP double-positive cells by fluorescent microscopy (Nikon ECLIPSE TE2000-E Inverted, Nikon, Melville, NY, USA; Zeiss Axio Observer Z1 motorized, Carl Zeiss Microscopy, Jena, Germany) and flow cytometry (Calibur or Canto II; BD Biosciences, Franklin Lakes, New Jersey, USA). Cells double positive for GFP and either mCherry or DR stain were live-cell sorted (AriaII BD; Biosciences) and re-plated for further analysis. The gating strategy used to flow-sort these cells is shown in **Supplementary Figure 1**.

### HIV-1 infected RTE cells co-culture with uninfected T-cells or monocytes

Flow-sorted GFP/mCherry double positive renal cells were co-cultured with the CD4+ T-cell line CEM-SS^[16]^ or the THP-1 monocytic cell line, at a ratio of 20:1 infected renal cells to uninfected T-cells or monocytes for 4 days, after which T-cells or monocytes were moved to a separate well. GFP expression was monitored over time by microscopy and flow cytometry.

### Drug Inhibition Studies

For the azidothymidine (AZT) and raltegravir inhibition studies, target RTE cells were pretreated with 100 and 10 μM of drug, respectively, at 37°C for 1 hour before co-culture with HIV-1 infected macrophages at 37°C still in the presence of the drugs. Drug treatment and co-culture were initiated 24-hours after macrophage infection. AZT and raltegravir were replenished daily and maintained in the co-culture media for the remainder of the experiment.

### Live imaging

Macrophage/RTE cell co-cultures were live imaged at 37°C using a Zeiss Axio Observer inverted microscope with stage incubator, CO2 buffering and an outer environmental chamber for up to 3 days. Images were taken at intervals of 4-6 minutes, and captured with a QUANTEM EMCCD camera (Photometrics, Tucson AZ) controlled by Metamorph (Molecular Devices, San Jose, CA). Images were analyzed and assembled using Metamorph software (Molecular Devices, San Jose, CA).

### Statistical Analyses

Each experiment was performed at least 3 times using PBMC and autologous RTE cells from 3 healthy donors. The interstitial inflammatory infiltrates of HIVAN in MICSS stained samples from **Figure1** are representative of many fields of view from one of the five HIVAN diagnosed biopsies. The frequency of the different cell types was assessed in all 5 biopsies and the results are shown in **figure 1** legend and the corresponding result section.

## RESULTS

### Macrophages are abundant in HIVAN inflammatory infiltrate

Renal macrophages are critical to homeostatic regulation of the kidney environment and their number increases following renal injury^[22]^. In the context of HIV-1 infection, the presence of either infected or uninfected macrophages in the kidney has the potential to fuel the dissemination of the virus within the renal tissue. To assess the role of macrophages in the spread of HIV in the kidney, we first evaluated their presence in the tissue lesions characteristic of HIVAN. We performed a multiplexed immunophenotyping analysis on 5 kidney biopsies from HIV-1 positive subjects diagnosed with HIVAN to examine the presence of macrophages in the inflammatory infiltrate. The mononuclear infiltrate was characterized by serially staining the tissue with markers for macrophages (CD68), T-cells (CD3), cytotoxic lymphocytes (CD8), B cells (CD20) and neutrophils (CD66b). We observed that macrophages constitute a high proportion of the inflammatory infiltrate within the HIVAN lesion and therefore a potential vehicle for viral spread in the kidney (**Fig. 1**). In the shown biopsy the number of positive cells/mm^2^ for each of the analyzed marker was as follow: 499 for CD20, 1102 for CD3, 998 for CD66b, 208 for CD68 and 544 for CD8. The average CD68 count for the 5 HIVAN biopsies analyzed was 626 cells/mm^2^ (median = 770, standard deviation 261), while the average CD3 count was 1028 (median = 1102, standard deviation 631). The CD68/CD3 ratio was inconsistent from case to case (in the example shown and in one other the number or T cells was higher; in a third case more macrophages were present, and the remaining two had about equal proportion of T cells and macrophages). Macrophages were present in 5/5 biopsies examined.

### Macrophages transfer HIV-1 to RTE cells

Prior work demonstrated that RTE cells are susceptible to productive HIV-1 infection following cell-cell interaction and virus transfer from infected T-cells^[16, 17]^. Given the consistent presence of macrophages in HIVAN biopsies (**Fig.1**), we hypothesized that macrophages could also contribute to infection of the renal epithelium. As macrophages are preferentially infected by CCR5-tropic strains of HIV-1^[36]^, we first evaluated the ability of different CCR5-tropic infectious molecular clones (IMC) of HIV-1 to infect macrophages and then assessed their ability to promote cell-to-cell infection of RTE cells. We used 3 previously described CCR5 tropic IMC of HIV-1 expressing GFP, HIV-ADA-GFP^[30]^, HIV-JRFL-GFP^[17]^, and HIV-sf162_EGFP^[31]^, to infect fully differentiated primary macrophages. We observed that the HIV-ADA-GFP and HIV-JRFL-GFP IMCs yielded the highest macrophage infection rates (19.4% and 27.1% respectively) (**Fig.2a**). We next tested the ability of these IMCs to directly infect RTE cells by incubating HPT-1b-mCherry cells with ≥15 MOI of each IMC. RTE cell infection was evaluated between day 3 and 7 post-infection by fluorescence microscopy. No RTE cells were found to express GFP in cultures where HPT-1b-mCherry were exposed to cell-free virus in the absence of macrophages or when a transwell membrane was used to impede contact between the two cell types (**Fig.2b** and **supplementary figure2**). To evaluate the ability of macrophages to mediate cell-to-cell infection of these HIV-1 IMCs to RTE cells, macrophages were differentiated in culture from primary monocytes for 6 days in the presence of MCSF and then co-cultured with the RTE cell line HPT-1b-mCherry (constitutively expressing mCherry). Twenty-four hours later, cells were incubated with either HIV-ADA-GFP, HIV-JRFL-GFP, or HIV-sf162-GFP IMCs for 8 hours after which cells were washed twice with PBS and incubated at 37°C for 6 additional days. Macrophages infected with the HIV-ADA-GFP were found to yield the highest cell-to-cell infection rate (from 2.7 to 5.6%) in culture (**Fig.2c**), therefore we selected this IMC for subsequent co-culture experiments. We observed higher cell death rates among cells infected with either the sf162 or the JRFL viruses compared to cells infected with the ADA clone. The higher toxicity observed for those viruses might explain why, despite demonstrating higher infection rates of macrophages, a lower number of measurable infected renal cells could be detected.

**Figure 2.**
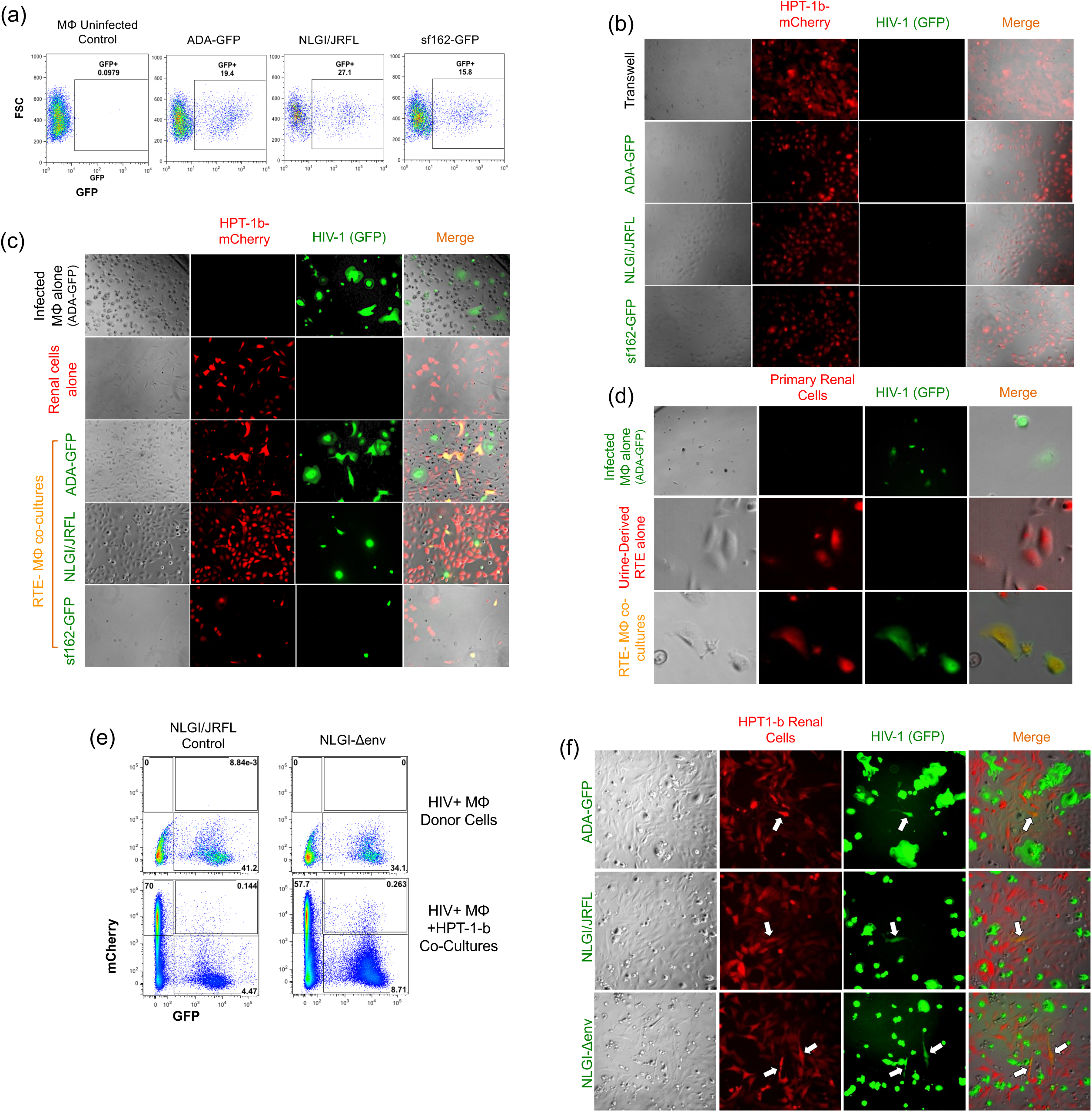
HIV-1 Infected macrophages mediate cell-to-cell infection when cocultured with renal tubule epithelial cells. **(a)** Monocyte derived macrophages were differentiated in presence of 20 ng/mL of MCSF for 7 days and then infected with 2.5 MOI of each of the indicated GFP-expressing IMCs. Infection rates were evaluated by assessing the percentage of GFP+ cells by flow-cytometry 7 days post infection. **(b)** HPT-1b-mCherry cells were incubated with ≥15 MOI of cell-free virus or separated from infected macrophages by a transwell membrane. No GFP expression could be detected in those conditions. HIV infected macrophages were co-cultured with either the HPT-1b-mCherry renal epithelial cell line **(c)** or with urine-derived autologous renal cells previously stained with the CellTracker Deep Red dye **(d)**. Transfer of virus from macrophages to HPT-1b-mCherry and primary renal cells was observed 3 days post-coculture as demonstrated by the presence of GFP/mCherry double positive cells. **(e-f)** Primary macrophages were infected with NLGI/JRFL or the JRFL pseudotyped envelope mutant virus NLGIΔenv for 24 hours prior to coculture with HPT-1b-mCherry renal cells. Infection of renal epithelial cells was evaluated by flow cytometry or fluorescence microscopy 6 days post-coculture with infected macrophages. White arrows indicate mCherry/GFP double positive renal cells.

To determine if primary RTE cells could acquire HIV-1 following co-culture with infected macrophages, we performed autologous co-cultures using RTE cells isolated directly from urine. As shown in **Figure2d and supplementary figure 4**, cell-to-cell infection of primary RTE cells was observed when co-cultured with autologous HIV-ADA-GFP infected macrophages. This demonstrates that direct contact between HIV-1 infected macrophages and RTE cells is required for renal epithelial cell infection.

To determine whether cell-to-cell infection from infected macrophages to renal cells requires the HIV-1 envelope glycoprotein, macrophages were infected with an envelope deficient HIV-1 IMC, NLGI-Δenv^[33]^, pseudotyped in trans with the JR-FL envelope to support a single round of viral entry into macrophages. To ensure that RTE cells were only exposed to the envelope deficient virus, co-cultures were initiated 24 hours post-infection, after extensive washing of infected macrophages with PBS. As a control, co-cocultures were conducted with macrophages infected with the unmutated version of this construct encoding a functional JRFL envelope. Cell-to-cell infection was detected in both control and mutant cocultures (**Fig.2e-f)**, demonstrating that virus transfer from macrophage to RTE cells does not involve HIV-1 envelope.

### HIV-1 infected renal tubule epithelial cells produce infectious virus as a result of viral transcription and integration

To further characterize the double positive population, mCherry+/GFP+ RTE cells derived from co-culture with infected macrophages were isolated by flow cytometry sorting as shown in **Fig.3a-b**, re-plated and examined by fluorescence microscopy. As noted above, the mCherry marker was stably introduced into RTE cells and serves as a functional marker of RTE cells, while GFP expression by those cells indicates HIV-1 infection. At 24 hours post-sort all the RTE cells isolated expressed both mCherry and GFP (**Fig.3c**), however at later time points, starting at day 3 post-sort, we observed that a portion of the cultured mCherry positive RTE cells no longer expressed GFP (**Fig.3d-e and Supplementary figure 3**), suggesting that in those cells the viral promoter became transcriptionally silent as seen in the latent state of HIV-1. By day 7 post-sort, about half of the cells present in the culture expressed only the mCherry marker (Fig. 3e).

**Figure 3.**
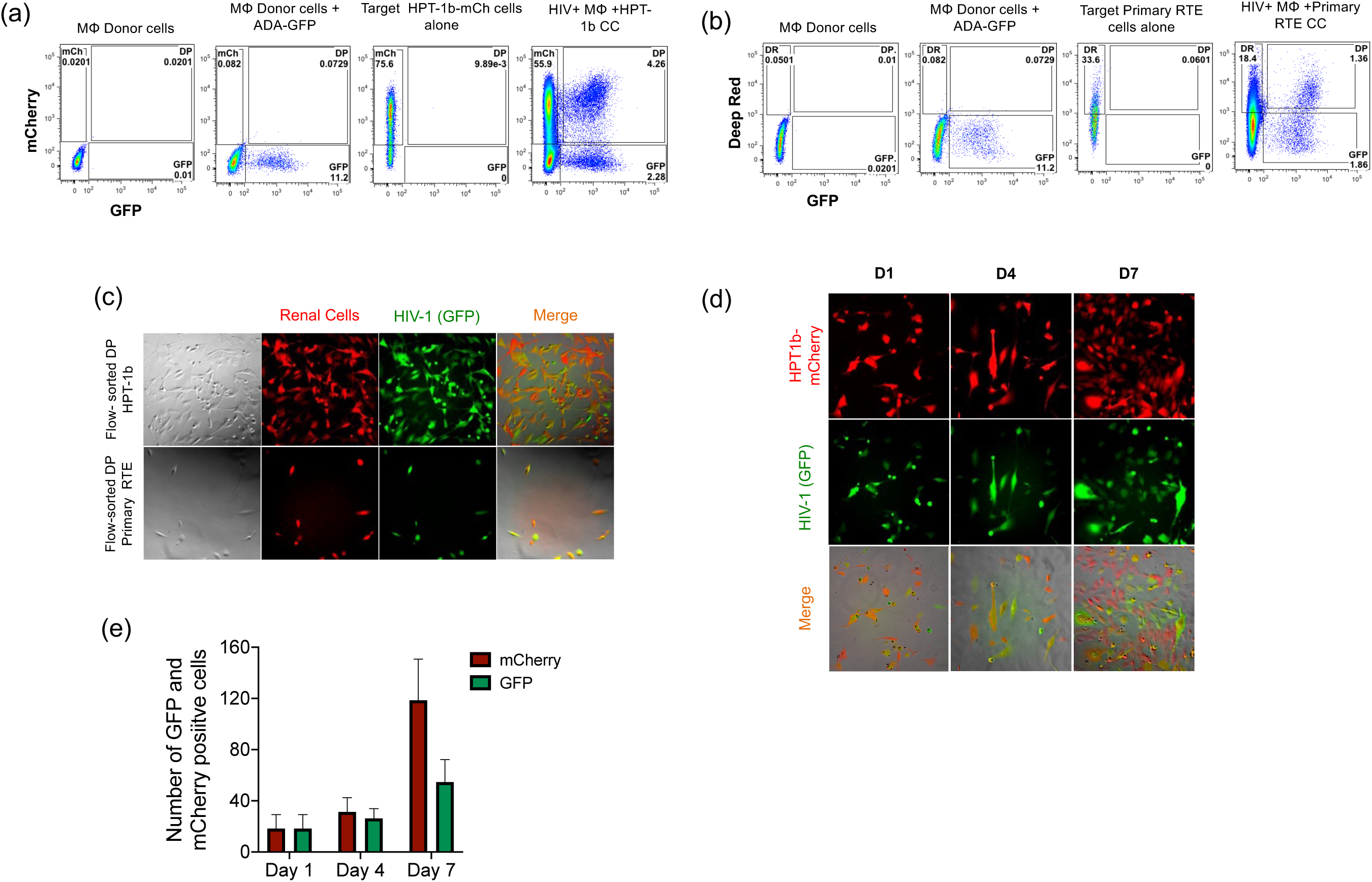
HIV gene expression in renal epithelial cells post co-culture with infected macrophages. Renal epithelial cells were plated with macrophages 24 hours prior to infection with 10 MOI of the HIV-ADA-GFP IMC, and co-cultured for 6 additional days. mCherry/GFP double positive HPT-1b **(a)** or urine-derived primary renal cells **(b)** were flow-sorted 7 days post-infection and re-plated for further analysis. **(c)** Flow-sorted cells were analyzed by fluorescence microscopy 24 hours post-sort to confirm the isolation of a pure population of mCherry/GFP double positive renal epithelial cells. **(d)** Time course microscopy analysis of HIV-GFP expression in flow-sorted mCherry/GFP double positive renal epithelial cells (days 1-7). **(e)** Number of cells, originally sorted as mCherry/GFP double positive, expressing GFP and/or mCherry at day 1, 4 and 7 post-sort. Data are shown as mean + standard deviation of the number of cells positive for each marker in 3 different fields.

To determine if the GFP expression observed in the RTE cells was the result of productive HIV-1 infection, co-cultures were carried out in presence of the reverse transcriptase inhibitor AZT, or the integrase inhibitor Raltegravir. HPT-1b-mCherry cells were treated with AZT or Raltegravir before and during co-culture with HIV-1 infected macrophages. Compared to the untreated co-cultures, we observed a 5-fold reduction in the percentage of mCherry/GFP double positive RTE cells in presence of AZT and a 12-fold reduction in presence of Raltegravir (**Fig.4a**), indicating that the GFP expression in this population required both HIV-1 reverse transcription and integration.

**Figure 4.**
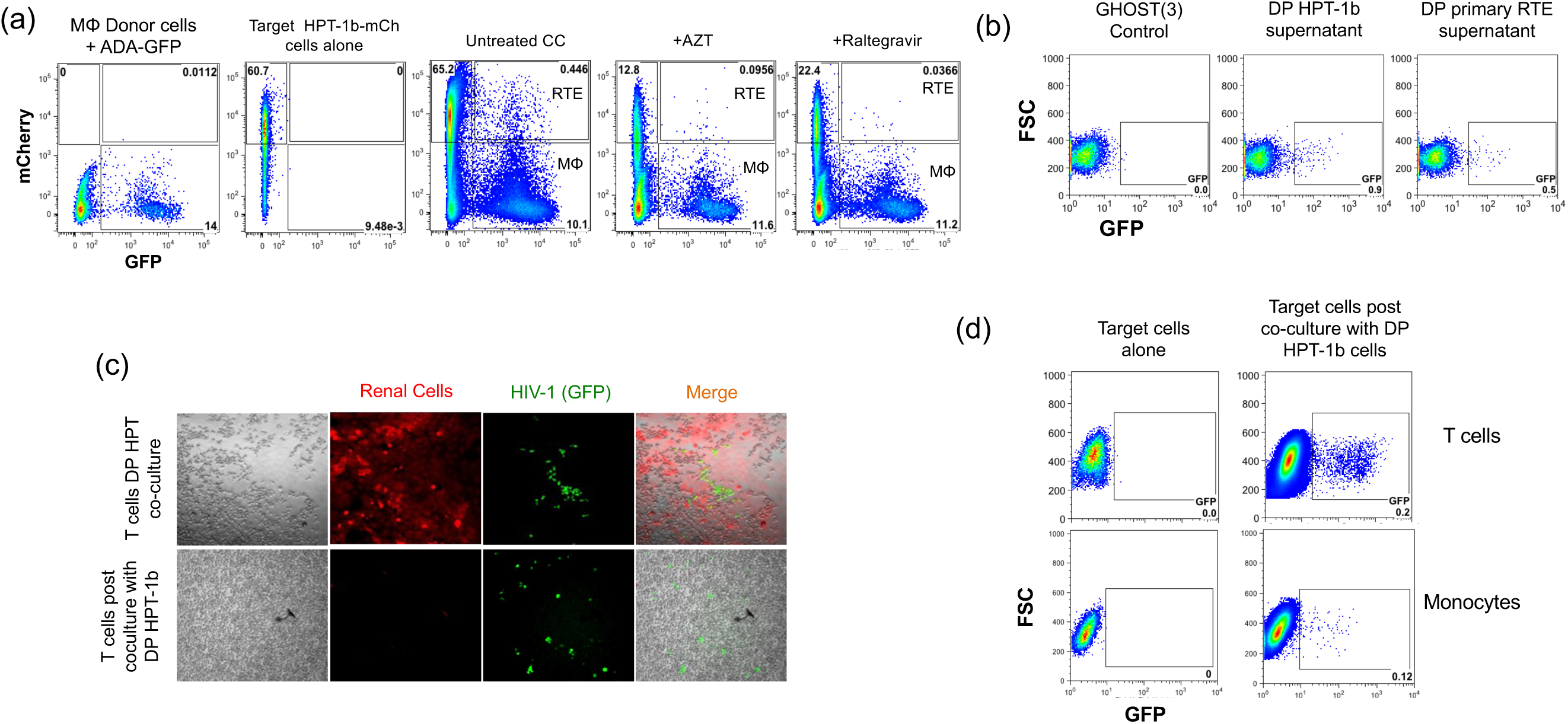
Infected renal cells produce infectious virus and transfer HIV-1 to target immune cells. **(a)** Primary macrophages were infected with HIV-GFP for 24 hours prior to co-culture with HPT-1b-mCherry renal cells. HPT-1b-mCherry renal cells were treated with AZT (100uM) or Raltegravir (10uM) for 1 hour prior to co-culture with infected macrophages and drugs were replenished every day throughout the remainder of the co-culture experiments. Infection of renal epithelial cells (mCherry/GFP+ double positive cells) in the coculture was evaluate after 6 days by flow-cytometry. **(b)** The GHOST(3) CXCR4+/CCR5+ indicator cell line was incubated with supernatants collected from flow-sorted mCherry/GFP double positive HPT-1b cells or primary urine-derived renal cells 4 days post-sort; the presence of infectious virions was evaluated by assessing the percentage of GFP+ cells by flow-cytometry 48 hours post infection. **(c)** CEM-SS T-cells or THP-1 monocytes were incubated for 7 days with mCherry/GFP double positive HPT-1b cells that had been in culture for 5 days after being flow-sorted. Transfer of virus from renal cells to T-cells or monocytes was evaluated by fluorescence microscopy **(c)** or flow-cytometry **(d)**.

To further confirm that these mCherry+/GFP+ renal cells were productively infected, we collected supernatants from flow-cytometry sorted GFP/mCherry double positive HPT-1b-mCherry or primary renal cells and assessed the presence of infectious virus in these supernatants using the GHOST(3) CXCR4+/CCR5+ reporter cell line. Supernatants collected at day 4 post-co-culture induced GFP expression in GHOST cells (**Fig.4b**), demonstrating production of infectious virus by mCherry/GFP double positive RTE cells. We next evaluated the ability of the flow sorted mCherry/GFP double positive RTE cells, to transfer HIV-1 to T-cells and monocytes using a co-culture system. As shown in **Fig.4c-d**, following co-culture of uninfected T-cells and monocytes with mCherry/GFP double positive renal cells both T-cells and monocytes became infected. These results demonstrate that following acquisition of HIV-1 from macrophages, renal tubule epithelial cells become productively infected with HIV-1 and can release virus that is infectious to cells that express CD4. These observations further validate and expand our previous findings and show bi-directional passage of HIV-1 between inflammatory cells, including T-cells and macrophages, that can be found in the renal interstitium, and renal epithelial cells^[16]^, thus defining a mechanism to sustain HIV-1 infection within the renal compartment.

### Live imaging of HIV-1 infected renal tubule epithelial cells reveals different cell fates

To explore the fate of HIV-1 infected RTE cells we performed daily monitoring of mCherry/GFP double positive RTE cells that acquired HIV-1 following co-culture with infected macrophages. We observed that in some fields the number of mCherry/GFP double positive cells increased in clusters over time (**Fig.5a**), suggesting either the occurrence of new viral transfer events or proliferation of infected renal cells. This observation is particularly interesting in light of the unique pattern of renal tubule infection observed in human kidney biopsies where HIV-1 infection was present in all the cells lining a single tubule, but was absent in neighboring tubules^[9, 13]^. To address this further we performed live imaging of HPT-1b-mCherry co-cultures from day 5 to 7 post infection. We observed multiple rounds of division by mCherry/GFP double positive renal cells leading to the formation of clusters of infected cells (**Fig.5b, Supplemental Movie1**). Division of HIV-1 infected primary renal epithelial cells was similarly observed in autologous co-cultures (**Fig.5c, Supplemental Movie2**). The daughter cells resulting from the observed cell divisions continued to express GFP, suggesting that the HIV-encoded GFP detected in double positive cells is a result of HIV-1 integration and production of virally encoded proteins. Given the close proximity of these clusters of double positive cells to the original infected cells in culture, it is likely this represents clonal proliferation of infected cells. To address this further, we sorted the mCherry/GFP double positive cells as single cells into 96 well plates (1 cell per well) and followed each single cell over time by fluorescence microscopy. As shown in **Fig.5d**, between day 2 and day 7 post-sort the number of mCherry/GFP double positive cells present in each well was higher than 1, consistent with clonal expansion of infected renal cells. Out of 510 wells each containing a single cell, we observed proliferation in 18 of them. We note that single cells grow poorly alone, so these numbers may underestimate this phenomenon. Proliferation of double positive cells also confirms their identity as renal cells, as macrophages are terminally differentiated cells that do not divide. In addition to the expansion of infected cells, two additional scenarios were observed in these cultures: infected renal cells became hypertrophic while continuing to express GFP (**Fig.5b, Supplemental Movie 1**) or died (**Fig.5d, Supplemental Movie 3**). These additional fates are consistent with primary pathological phenotypes reported in HIVAN biopsies^[7, 37]^. As noted above, some of the double positive sorted cells lost GFP expression over time, consistent with the promoter silencing seen in latently infected cells; suggesting latency as a fourth fate for infected RTE.

**Figure 5.**
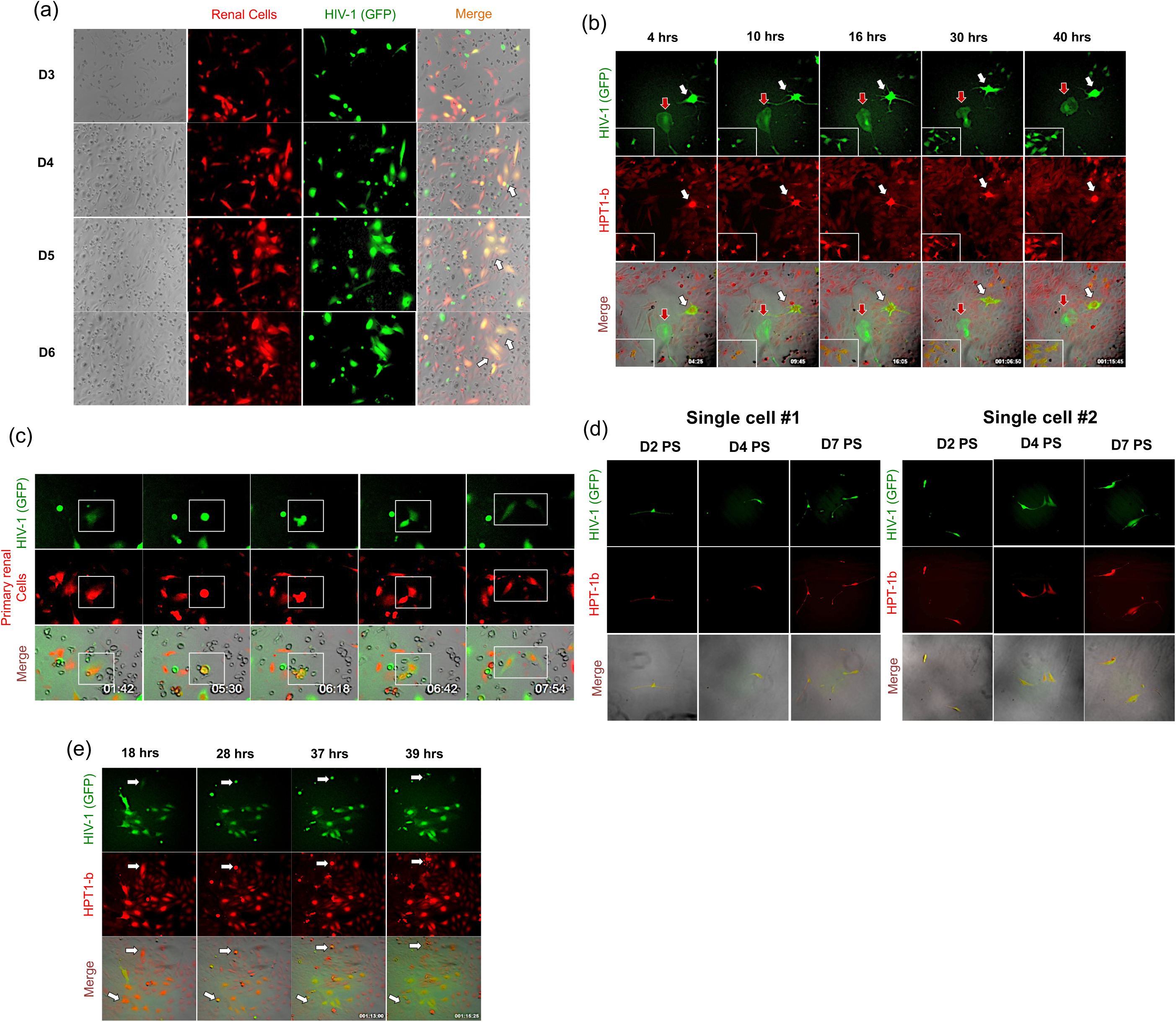
Proliferation of HIV-1 infected renal epithelial cells. Macrophage HPT-1b-mCherry co-cultures were analyzed over time by fluorescence microscopy to assess viral transfer and the fate of infected renal cells. Clusters of mCherry/GFP double positive renal cells began to appear at day 4 post-coculture (white arrows in Merge images in **panel a**). Live imaging of cocultures between HIV infected macrophages and either HPT-1b-mCherry **(b)** or primary urine-derived renal epithelial cells **(c)** between days 5 and 7 post-infection. Imaging demonstrates three different cellular fates: 1. Clustered proliferation of infected mCherry/GFP double positive renal cells (**panels b, c and d**); 2. Hypertrophy, persistent GFP expression and no cellular division (white arrows in **panel b**); or 3. cell death (white arrows in **panel e**). Red arrows in b indicate an HIV infected macrophage. Numbers indicate elapsed time since beginning of live imaging. White boxes in b and c delimit areas where cell proliferation was observed (see supplemental movies 1 and 2).

## DISCUSSION

This study demonstrates that HIV-1 infected macrophages mediate cell-to-cell infection of RTE cells, elucidating an additional mechanism that the virus could exploits to infect and spread in the renal epithelium. As demonstrated using multiplexed immunohistochemistry analysis of kidney biopsies from four HIV-1 positive individuals with HIVAN, macrophages are abundant within the inflammatory infiltrate of HIVAN lesions. The recruitment/presence of HIV-1 infected macrophages into the kidney during the tissue-damage response can therefore facilitate the spread of the virus to neighboring renal epithelial cells. Both primary urine-derived RTE cells or the RTE cell line HPT-1b-mCherry, acquire HIV-1 following co-culture with infected macrophages. Virus infection was not observed when the two cell types were physically separated by a transwell membrane, or when high MOI of cell-free virus was added directly to RTE cells, confirming that cell-cell contact is required for successful transfer of HIV-1. We show that RTE cells that acquired HIV-1 from infected macrophages produce infectious virus and transmit the virus back to lymphoid cells, including T-cells and monocytes. *In vivo* HIV-1 positive individuals shed virus in urine^[38]^ and those viruses are genetically different from the viral quasi-species found in blood but are closely related to urine-derived renal epithelial cells^[39]^, supporting renal epithelial cells as one source of urine viruses.

Interestingly we observed four cellular fates for HIV-1 infected RTE cells: hypertrophy, cell death, proliferation and transcriptional silencing, consistent with viral latency. We have previously reported the presence of hypertrophic tubule cells in the Tg26 mouse model of renal infection and in HIVAN human biopsies, and demonstrated that this phenotype, together with cell cycle arrest and polyploidy are primarily induced by the expression of HIV-1 *Vpr* and its ability to impair cytokinesis^[40-42]^. The recapitulation of all those phenotypes in the *in vitro* model described here demonstrates that HIV-1 gene expression is responsible for the phenotypic changes observed in renal biopsies from HIV-1 infected individuals that correlated with renal dysfunction^[5]^.

To our knowledge this is the first demonstration of proliferation of HIV-1 infected renal tubule cells consistent with clonal expansion. Renal biopsies from infected individuals examined by RNA situ hybridization demonstrate infection in circumferential neighboring cells in a single renal tubule interspersed with areas of uninfected tubules^[9, 13]^. This pattern could be accounted for by the clonal expansion of an infected renal epithelial cell in response to injury in the kidney^[43]^.

Clonal amplification has recently emerged as one of the mechanisms through which HIV-1 infected CD4+ T-cells persist and expand in ART treated individuals^[44, 45]^. Proviral DNA integrated near oncogenes is replicated along with host genetic material during cell division, increasing the pool of infected cells in a host^[46, 47]^. Our *in vitro* observation that infected RTE cells can clonally expand similarly to CD4+ T-cells, highlights the importance of considering the role of non-lymphoid viral compartments in HIV-1 persistence. Sequence analyses of the proviruses and/or their integration sites would support the possibility that cellular proliferation maintains the HIV reservoir in renal tubule cells. Future studies will also assess whether the HIV-1 integration sites in infected renal epithelial cells plays a role in their clonal expansion, as reported for CD4+ T-cells^[46-48]^.

In summary, we show that in addition to T-cells, macrophages can transfer HIV-1 to renal epithelial cells, and thus could contribute to viral spread within the kidney in light of the inflammatory interstitial infiltrate. These results support a scenario in which infected macrophages present in the renal tissue could initiate or perpetuate infection by transferring virus to renal epithelial cells, which in turn, can infect susceptible lymphoid cells. Furthermore, once infected, renal epithelial cells can undergo clonal proliferation, with or without transcriptional silencing, providing a mechanism for HIV-1 persistence in the kidney. As cure strategies advance, it will be important to understand the dynamics of viral persistence and expression in both lymphoid and non-lymphoid reservoirs.

---

**Supplementary figure 1.**
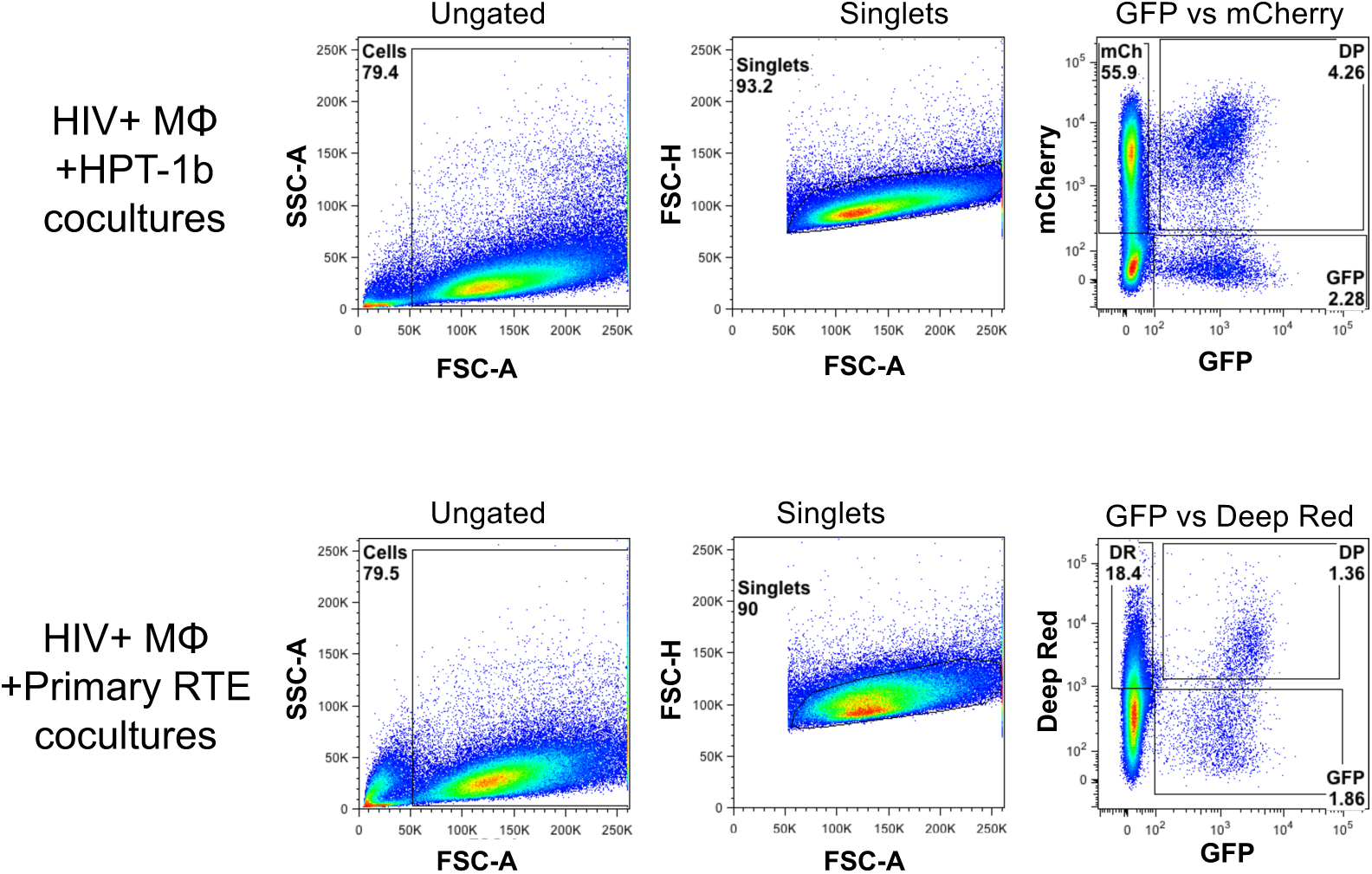
Gating strategy used to sort mCherry/GFP double positive renal cells.

**Supplementary figure 2.**
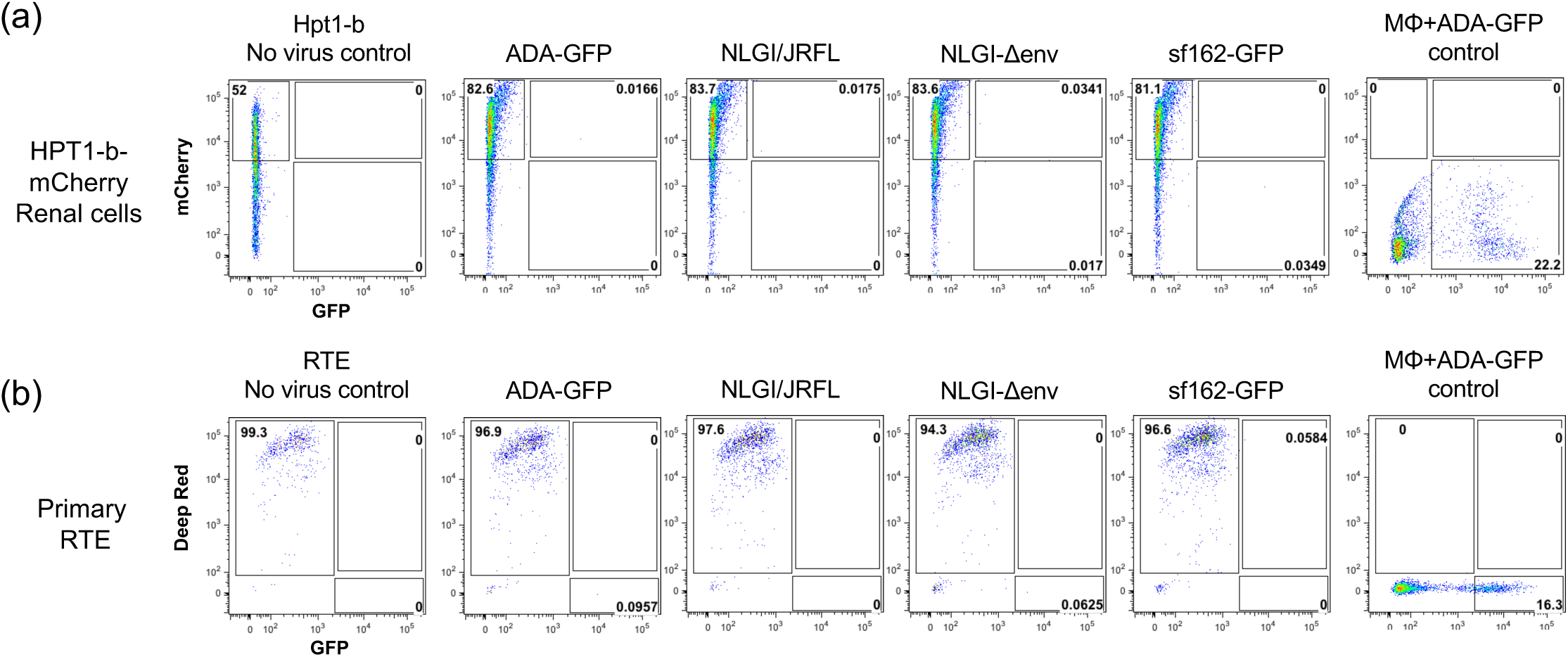
Renal epithelial cells are not susceptible to cell-free HIV-1 infection. HPT-1b-mCherry (**a**), DeepRed stained primary renal epithelial cells (**b**) were incubated with ≥15 MOI of each of the indicated HIV-1 IMCs for 8 hours, and then cultured for 6 days. As positive control, macrophage infection with 6 MOI of ADA-GFP was performed in parallel. HIV infection was evaluated by flow-cytometry 6 days post-virus incubation.

**Supplementary figure 3.**
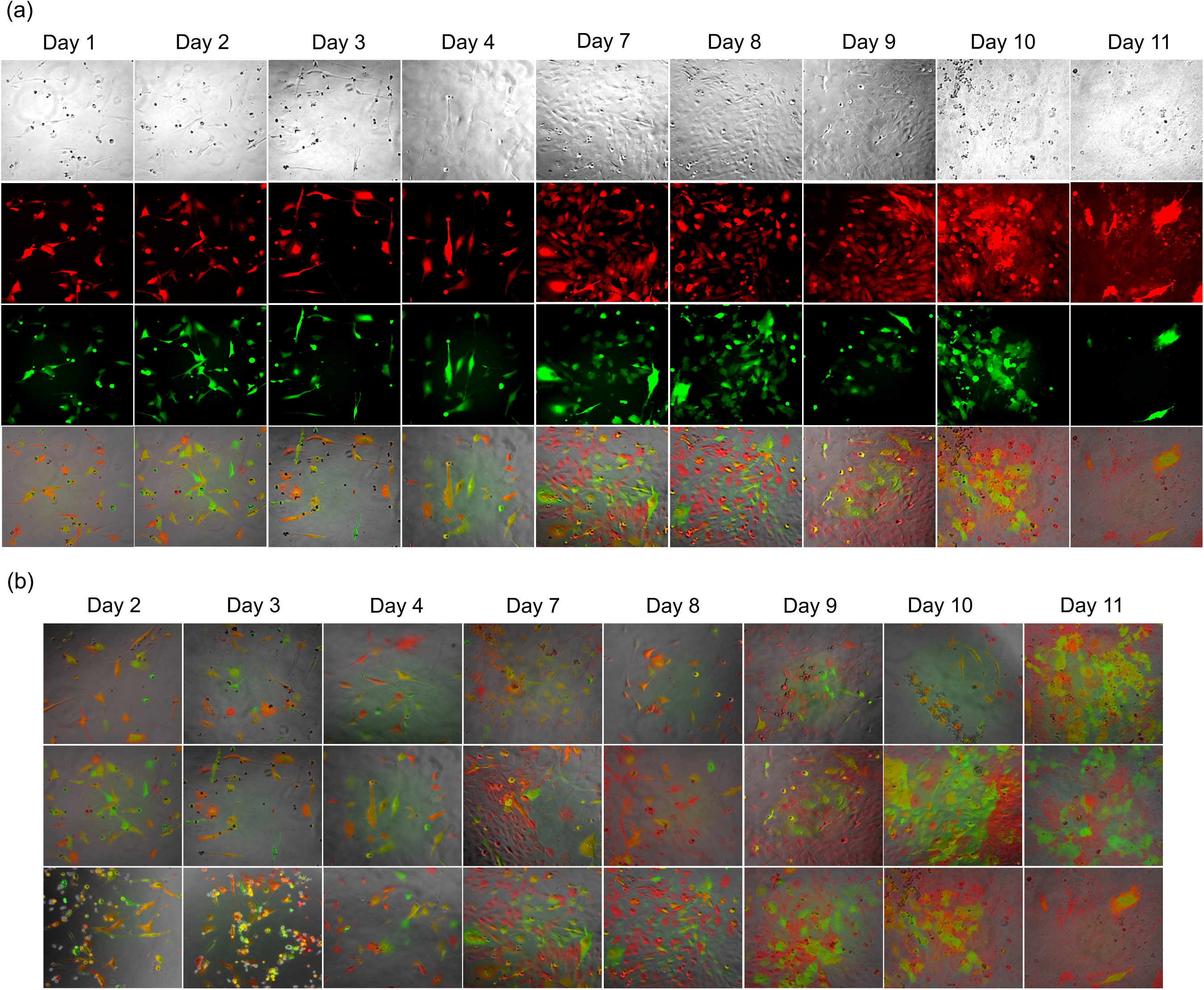
HIV gene expression in renal epithelial cells post co-culture with infected macrophages. mCherry/GFP double positive HPT-1b were flow-sorted 7 days post-coculture with HIV-infected macrophages and re-plated for further analysis. **(a)** Time course microscopy analysis of HIV-GFP expression in flow-sorted mCherry/GFP double positive renal epithelial cells between day 1 and 11 post-sort. Additional analyzed fields of view are shown in (**b**).

**Supplementary figure 4:**
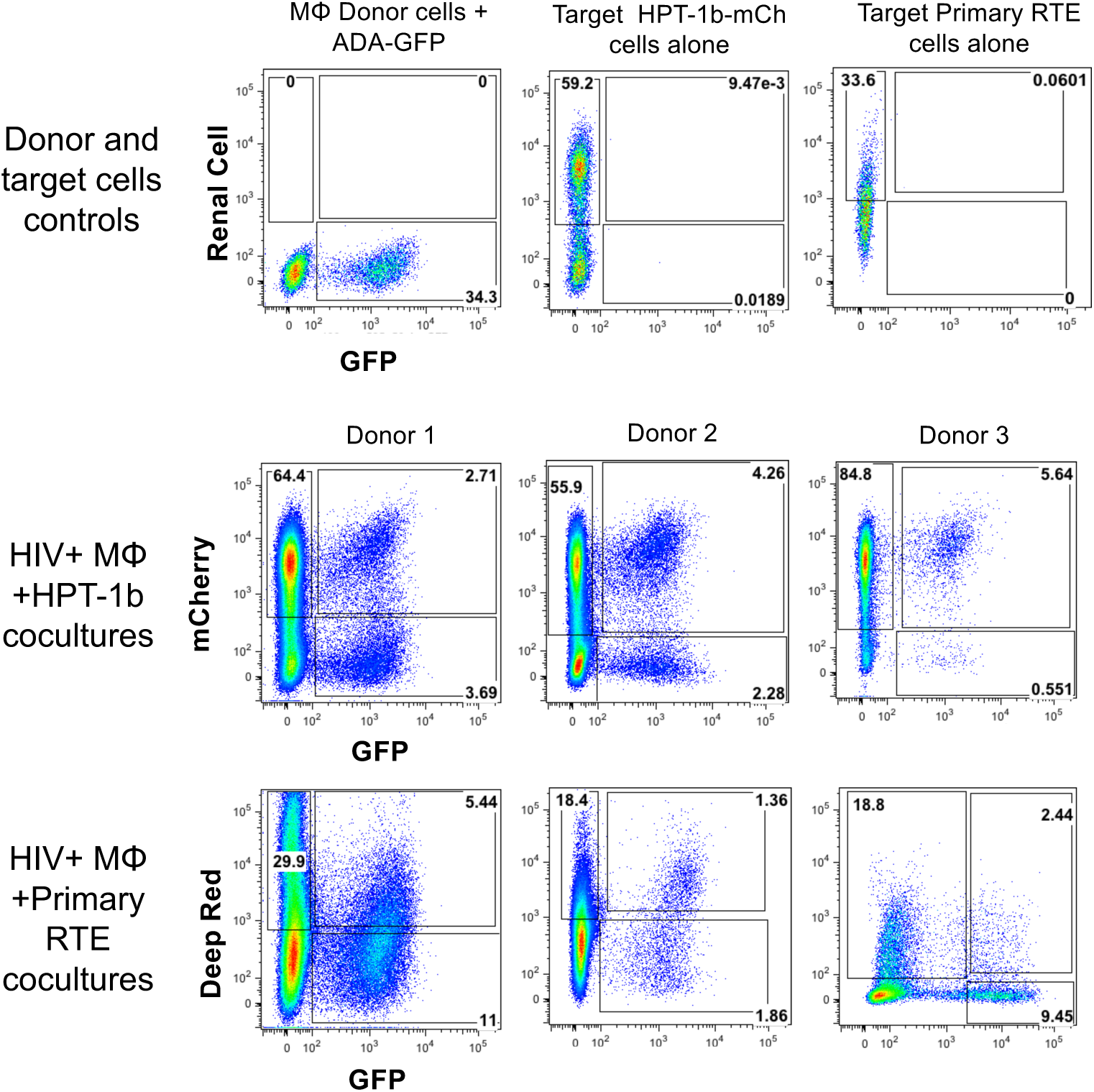
Renal epithelial cells were plated with macrophages 24 hours prior to infection with 10 MOI of the HIV-ADA-GFP IMC, and co-cultured for 6 additional days. mCherry/GFP double positive HPT-1b (middle row) or urine-derived primary renal cells (bottom row) were flow-sorted 7 days post-infection and re-plated for further analysis. Co-cultures were performed using macrophages and urine-derived primary renal cells from 3 different donors. Representative single cell controls are shown in the top row.

## Author Contributions

K.H performed all co-culture experiments and analyzed the data. A.G., S.G and B.C. performed and analyzed the multiparametric immunohistochemistry analysis. M.B and M.K. designed the study, oversaw experiments, analyzed results and edited the manuscript.

## Acknowledgements

This work was supported by the National Institute of Diabetes and Digestive and Kidney Diseases (NIDDK) grants number P01DK056492 and R01DK108367.

The authors thank Ilaria Laface for assistance with MICSSS and Christina Wyatt and Vivette D’Agati for providing the kidney biopsies.

The following reagents were obtained through the NIH AIDS Reagent Program, Division of AIDS, NIAID, NIH: Azidothymidine (AZT) and Raltegravir.

We thank the Duke Light Microscopy Core for the use of the shared instrumentation including the Zeiss Axio Observer Z1 microscope and Metamorph software, supported by NIH Shared Instrumentation grant 1S10OD020010-01A1.

## Conflict of Interests

None

## Supplementary material

- Supplemental Movie 1: Clonal expansion of mCherry/GFP DP HPT-1b cells and hypertrophy
- Supplemental Movie 2: Clonal expansion of mCherry/GFP DP Primary renal cells
- Supplemental Movie 3: Cell death of mCherry/GFP DP HPT-1b cells and hypertrophy

## References

1. Siliciano JD, Kajdas J, Finzi D, Quinn TC, Chadwick K, Margolick JB, et al. Long-term follow-up studies confirm the stability of the latent reservoir for HIV-1 in resting CD4+ T cells. Nat Med 2003; 9(6):727–728.

2. Boritz EA, Douek DC. Perspectives on Human Immunodeficiency Virus (HIV) Cure: HIV Persistence in Tissue. J Infect Dis 2017; 215(suppl_3):S128–S133.

3. Wong JK, Yukl SA. Tissue reservoirs of HIV. Curr Opin HIV AIDS 2016; 11(4):362–370.

4. Avettand-Fenoel V, Rouzioux C, Legendre C, Canaud G. HIV Infection in the Native and Allograft Kidney: Implications for Management, Diagnosis, and Transplantation. Transplantation 2017; 101(9):2003–2008.

5. Swanepoel CR, Atta MG, D’Agati VD, Estrella MM, Fogo AB, Naicker S, et al. Kidney disease in the setting of HIV infection: conclusions from a Kidney Disease: Improving Global Outcomes (KDIGO) Controversies Conference. Kidney Int 2018; 93(3):545–559.

6. Kopp JB, Klotman ME, Adler SH, Bruggeman LA, Dickie P, Marinos NJ, et al. Progressive glomerulosclerosis and enhanced renal accumulation of basement membrane components in mice transgenic for human immunodeficiency virus type 1 genes. Proc Natl Acad Sci U S A 1992; 89(5):1577–1581.

7. D’Agati V, Suh JI, Carbone L, Cheng JT, Appel G. Pathology of HIV-associated nephropathy: a detailed morphologic and comparative study. Kidney Int 1989; 35(6):1358–1370.

8. Kimmel PL, Ferreira-Centeno A, Farkas-Szallasi T, Abraham AA, Garrett CT. Viral DNA in microdissected renal biopsy tissue from HIV infected patients with nephrotic syndrome. Kidney Int 1993; 43(6):1347–1352.

9. Bruggeman LA, Ross MD, Tanji N, Cara A, Dikman S, Gordon RE, et al. Renal epithelium is a previously unrecognized site of HIV-1 infection. J Am Soc Nephrol 2000; 11(11):2079–2087.

10. Cohen AH, Sun NC, Shapshak P, Imagawa DT. Demonstration of human immunodeficiency virus in renal epithelium in HIV-associated nephropathy. Mod Pathol 1989; 2(2):125–128.

11. Marras D, Bruggeman LA, Gao F, Tanji N, Mansukhani MM, Cara A, et al. Replication and compartmentalization of HIV-1 in kidney epithelium of patients with HIV-associated nephropathy. Nat Med 2002; 8(5):522–526.

12. Wyatt CM, Meliambro K, Klotman PE. Recent progress in HIV-associated nephropathy. Annu Rev Med 2012; 63:147–159.

13. Canaud G, Dejucq-Rainsford N, Avettand-Fenoel V, Viard JP, Anglicheau D, Bienaime F, et al. The kidney as a reservoir for HIV-1 after renal transplantation. J Am Soc Nephrol 2014; 25(2):407–419.

14. Eitner F, Cui Y, Hudkins KL, Stokes MB, Segerer S, Mack M, et al. Chemokine receptor CCR5 and CXCR4 expression in HIV-associated kidney disease. J Am Soc Nephrol 2000; 11(5):856–867.

15. Hatsukari I, Singh P, Hitosugi N, Messmer D, Valderrama E, Teichberg S, et al. DEC-205-mediated internalization of HIV-1 results in the establishment of silent infection in renal tubular cells. J Am Soc Nephrol 2007; 18(3):780–787.

16. Blasi M, Balakumaran B, Chen P, Negri DR, Cara A, Chen BK, et al. Renal epithelial cells produce and spread HIV-1 via T-cell contact. AIDS 2014; 28(16):2345–2353.

17. Chen P, Chen BK, Mosoian A, Hays T, Ross MJ, Klotman PE, et al. Virological synapses allow HIV-1 uptake and gene expression in renal tubular epithelial cells. J Am Soc Nephrol 2011; 22(3):496–507.

18. Sattentau QJ, Stevenson M. Macrophages and HIV-1: An Unhealthy Constellation. Cell Host Microbe 2016; 19(3):304–310.

19. Castellano P, Prevedel L, Eugenin EA. HIV-infected macrophages and microglia that survive acute infection become viral reservoirs by a mechanism involving Bim. Sci Rep 2017; 7(1):12866.

20. Raposo G, Moore M, Innes D, Leijendekker R, Leigh-Brown A, Benaroch P, et al. Human macrophages accumulate HIV-1 particles in MHC II compartments. Traffic 2002; 3(10):718–729.

21. Tang PM, Nikolic-Paterson DJ, Lan HY. Macrophages: versatile players in renal inflammation and fibrosis. Nat Rev Nephrol 2019; 15(3):144–158.

22. Rogers NM, Ferenbach DA, Isenberg JS, Thomson AW, Hughes J. Dendritic cells and macrophages in the kidney: a spectrum of good and evil. Nat Rev Nephrol 2014; 10(11):625–643.

23. Remark R, Merghoub T, Grabe N, Litjens G, Damotte D, Wolchok JD, et al. In-depth tissue profiling using multiplexed immunohistochemical consecutive staining on single slide. Sci Immunol 2016; 1(1):aaf6925.

24. Akturk G, Sweeney R, Remark R, Merad M, Gnjatic S. Multiplexed Immunohistochemical Consecutive Staining on Single Slide (MICSSS): Multiplexed Chromogenic IHC Assay for High-Dimensional Tissue Analysis. Methods Mol Biol 2020; 2055:497–519.

25. Bankhead P, Loughrey MB, Fernandez JA, Dombrowski Y, McArt DG, Dunne PD, et al. QuPath: Open source software for digital pathology image analysis. Sci Rep 2017; 7(1):16878.

26. Ross MJ, Wosnitzer MS, Ross MD, Granelli B, Gusella GL, Husain M, et al. Role of ubiquitin-like protein FAT10 in epithelial apoptosis in renal disease. J Am Soc Nephrol 2006; 17(4):996–1004.

27. Blasi M, Negri D, LaBranche C, Alam SM, Baker EJ, Brunner EC, et al. IDLV-HIV-1 Env vaccination in non-human primates induces affinity maturation of antigen-specific memory B cells. Commun Biol 2018; 1:134.

28. Zhou T, Benda C, Dunzinger S, Huang Y, Ho JC, Yang J, et al. Generation of human induced pluripotent stem cells from urine samples. Nat Protoc 2012; 7(12):2080–2089.

29. Zhou T, Benda C, Duzinger S, Huang Y, Li X, Li Y, et al. Generation of induced pluripotent stem cells from urine. J Am Soc Nephrol 2011; 22(7):1221–1228.

30. Dave VP, Hajjar F, Dieng MM, Haddad E, Cohen EA. Efficient BST2 antagonism by Vpu is critical for early HIV-1 dissemination in humanized mice. Retrovirology 2013; 10:128.

31. Brown A, Gartner S, Kawano T, Benoit N, Cheng-Mayer C. HLA-A2 down-regulation on primary human macrophages infected with an M-tropic EGFP-tagged HIV-1 reporter virus. J Leukoc Biol 2005; 78(3):675–685.

32. Cohen GB, Gandhi RT, Davis DM, Mandelboim O, Chen BK, Strominger JL, et al. The selective downregulation of class I major histocompatibility complex proteins by HIV-1 protects HIV-infected cells from NK cells. Immunity 1999; 10(6):661–671.

33. Durham ND, Chen BK. HIV-1 Cell-Free and Cell-to-Cell Infections Are Differentially Regulated by Distinct Determinants in the Env gp41 Cytoplasmic Tail. J Virol 2015; 89(18):9324–9337.

34. Yang FC, Kuang WD, Li C, Sun WW, Qu D, Wang JH. Toll-Interacting Protein Suppresses HIV-1 Long-Terminal-Repeat-Driven Gene Expression and Silences the Post-Integrational Transcription of Viral Proviral DNA. PLoS One 2015; 10(4):e0125563.

35. Morner A, Bjorndal A, Albert J, Kewalramani VN, Littman DR, Inoue R, et al. Primary human immunodeficiency virus type 2 (HIV-2) isolates, like HIV-1 isolates, frequently use CCR5 but show promiscuity in coreceptor usage. J Virol 1999; 73(3):2343–2349.

36. Quitadamo B, Peters PJ, Repik A, O’Connell O, Mou Z, Koch M, et al. HIV-1 R5 Macrophage-Tropic Envelope Glycoprotein Trimers Bind CD4 with High Affinity, while the CD4 Binding Site on Non-macrophage-tropic, T-Tropic R5 Envelopes Is Occluded. J Virol 2018; 92(2).

37. Cohen AH, Nast CC. HIV-associated nephropathy. A unique combined glomerular, tubular, and interstitial lesion. Mod Pathol 1988; 1(2):87–97.

38. Blasi M, Carpenter JH, Balakumaran B, Cara A, Gao F, Klotman ME. Identification of HIV-1 genitourinary tract compartmentalization by analyzing the env gene sequences in urine. AIDS 2015; 29(13):1651–1657.

39. Blasi M, Stadtler H, Chang J, Hemmersbach-Miller M, Wyatt C, Klotman P, et al. Detection of Donor’s HIV Strain in HIV-Positive Kidney-Transplant Recipient. N Engl J Med 2020; 382(2):195–197.

40. Dickie P, Roberts A, Uwiera R, Witmer J, Sharma K, Kopp JB. Focal glomerulosclerosis in proviral and c-fms transgenic mice links Vpr expression to HIV-associated nephropathy. Virology 2004; 322(1):69–81.

41. Payne EH, Ramalingam D, Fox DT, Klotman ME. Polyploidy and Mitotic Cell Death Are Two Distinct HIV-1 Vpr-Driven Outcomes in Renal Tubule Epithelial Cells. J Virol 2018; 92(2).

42. Rosenstiel PE, Gruosso T, Letourneau AM, Chan JJ, LeBlanc A, Husain M, et al. HIV-1 Vpr inhibits cytokinesis in human proximal tubule cells. Kidney Int 2008; 74(8):1049–1058.

43. Kusaba T, Lalli M, Kramann R, Kobayashi A, Humphreys BD. Differentiated kidney epithelial cells repair injured proximal tubule. Proc Natl Acad Sci U S A 2014; 111(4):1527–1532.

44. Reeves DB, Duke ER, Wagner TA, Palmer SE, Spivak AM, Schiffer JT. A majority of HIV persistence during antiretroviral therapy is due to infected cell proliferation. Nat Commun 2018; 9(1):4811.

45. Coffin JM, Wells DW, Zerbato JM, Kuruc JD, Guo S, Luke BT, et al. Clones of infected cells arise early in HIV-infected individuals. JCI Insight 2019; 4(12).

46. Maldarelli F, Wu X, Su L, Simonetti FR, Shao W, Hill S, et al. HIV latency. Specific HIV integration sites are linked to clonal expansion and persistence of infected cells. Science 2014; 345(6193):179–183.

47. Wagner TA, McLaughlin S, Garg K, Cheung CY, Larsen BB, Styrchak S, et al. HIV latency. Proliferation of cells with HIV integrated into cancer genes contributes to persistent infection. Science 2014; 345(6196):570–573.

48. Haworth KG, Schefter LE, Norgaard ZK, Ironside C, Adair JE, Kiem HP. HIV infection results in clonal expansions containing integrations within pathogenesis-related biological pathways. JCI Insight 2018; 3(13).

